# BRG1 promotes transcriptional patterns that are permissive to proliferation in cancer cells

**DOI:** 10.1101/2020.07.03.187385

**Authors:** Katherine A. Giles, Cathryn M. Gould, Joanna Achinger-Kawecka, Scott G. Page, Georgia Kafer, Phuc-Loi Luu, Anthony J. Cesare, Susan J. Clark, Phillippa C. Taberlay

## Abstract

**Background:** BRG1 (encoded by *SMARCA4*) is a catalytic component of the SWI/SNF chromatin remodelling complex, with key roles in modulating DNA accessibility. Dysregulation of BRG1 is observed, but functionally uncharacterised, in a wide range of malignancies. We have probed the functions of BRG1 on a background of prostate cancer to investigate how BRG1 controls gene expression programs and cancer cell behaviour.

**Results:** Our investigation of *SMARCA4* revealed that BRG1 is universally overexpressed in 486 tumours from The Cancer Genome Atlas prostate cohort, as well as in a complementary panel of 21 prostate cell lines. Next, we utilised a temporal model of BRG1 depletion to investigate the molecular effects on global transcription programs. Unexpectedly, depleting BRG1 had no impact on alternative splicing and conferred only modest effect on global expression. However, of the transcriptional changes that occurred, most manifested as down-regulated expression. Deeper examination found the common thread linking down-regulated genes was involvement in proliferation, including several known to increase prostate cancer proliferation (*KLK2*, *PCAT1* and *VAV3*). Interestingly, the promoters of genes driving proliferation were bound by BRG1 as well as the oncogenic transcription factors, AR and FOXA1. We also noted that BRG1 depletion repressed genes involved in cell cycle progression and DNA replication but intriguingly, these pathways operated independently of AR and FOXA1. In agreement with transcriptional changes, depleting BRG1 conferred G1 arrest.

**Conclusions:** Our data have revealed that BRG1 has capacity to drive oncogenesis by coordinating oncogenic pathways dependent on BRG1 for proliferation, cell cycle progression and DNA replication.

## BACKGROUND

Nucleosomes serve as a physical backbone for chromatin organization on a global scale and at local gene regulatory elements. Nucleosomes therefore govern both genome-wide stability and local DNA accessibility (1). Nucleosome positioning by ATP-dependent chromatin remodellers plays a critical role in regulating DNA accessibility and allows genes to be expressed at the appropriate place and time (1). Genomic profiling has demonstrated that dynamic regulation of DNA accessibility occurs primarily at DNA regulatory elements, which are cell type specific, and that DNA accessibility changes reflect concomitant transcriptional patterns. (2, 3). It is essential for chromatin to be relaxed at active gene promoters to create an ordered nucleosome disassembly, which permits binding of RNA pol II and the general transcription machinery (4, 5). In agreement, ChIP-seq data show that transcription factors are concentrated on accessible DNA, with the highest levels of bound transcription factors correlating with the most accessible genomic regions (6). Conversely, chromatin condensation resulting in reduced DNA accessibility is necessary for transcriptional repression (7). Disruption to the DNA accessibility landscape is a feature of cancer (2, 8, 9). This was recently emphasized in genomic sequencing data from multiple cancers and cancer subtypes, which revealed associations between the accessible chromatin organization and mutation load (8). Moreover, studies of aged human and yeast cells demonstrated that nucleosome loss compromises genome stability, gene regulation and transcription (10, 11).

Genes encoding ATP-dependent chromatin remodellers are themselves frequently mutated and often atypically expressed in cancer (5, 12–16). Notably, the SWI/SNF chromatin remodelling complex is mutated or transcriptionally deregulated in ~20% of cancers; a mutation frequency approaching that of *TP53* (~26%) (12, 14, 17). The SWI/SNF complex is often described as a tumour suppressor because it is required by the Retinoblastoma protein (Rb) family for regulation of normal cell growth (18, 19). Disruptions of multiple SWI/SNF subunits are reported in human tumours and cell lines (13–15, 20–37), often accompanied by a loss of heterozygosity consistent with the inactivation of a tumour suppressor (13, 34). The specific SWI/SNF mutations observed in tumours and the cancers associated with altered SWI/SNF function have been extensively reviewed (12–15, 26, 31, 34, 38). However, the mechanism and functional consequences of SWI/SNF dysregulation are still being defined.

Brahma-related gene 1 (BRG1) is one of the two mutually exclusive ATPases within the SWI/SNF complex. Interestingly, *SMARCA4*, the gene encoding BRG1, has been observed in both down- and up-regulated states in cancer, indicative of the diverse and complex BRG1 functions. *SMARCA4* mRNA was seen to be down regulated in bladder, colon, non-triple negative breast cancers, head and neck, oesophageal, melanoma, pancreatic, lung and ovarian cancers, and *SMARCA4* mutation rates in these cancers have been reported between 4-13% (12–14, 22, 24, 30, 39–41). In contrast, *SMARCA4* has been reported as over expressed in cancers of the prostate, triple negative breast cancers and some leukaemias (12, 22, 24, 30, 42, 43). In *SMARCA4* over expressing cancers, no significant recurrent mutations have been reported (42, 44–46). The importance of BRG1 in cancer is further evidenced through studies of synthetic lethality, where BRG1 was observed to have a synthetic lethal relationship with the alternative SWI/SNF ATPase Braham (BRM), and Aurora A kinase in lung cancer, and PTEN in prostate cancer (43, 47, 48).

Examination of multiple prostate cancer cohorts has demonstrated elevated *SMARCA4* expression or increased BRG1 protein levels. Clinical studies of primary prostate tumours reported an overall increase in BRG1 protein by immunohistochemistry (42, 44–46). Moreover, increased *SMARCA4* gene expression has been reported in tumours from The Cancer Genome Atlas (TCGA) prostate cancer cohort (49, 50). While it is established that BRG1 is commonly up regulated in prostate cancer, the full range of molecular pathways impacted by dysregulated BRG1 levels and the contribution of these molecular changes to the atypical phenotype of prostate cancer cells remains unclear.

BRG1 has known roles in regulating DNA for temporal gene expression at both promoters and enhancer gene regulatory elements (4, 51–56). Moreover, BRG1 maintains the epigenetic landscape of a cell at these gene regulatory elements. Specifically, BRG1 has been directly linked to transcriptional output through its recognition of H3K14ac (57–59). In the absence of H3K14ac, BRG1 is still present at promoters and histones are disassembled from the chromatin; however, transcription is reduced (60). At enhancers, BRG1 depletion greatly reduces H3K27ac and subtlety reduces H3K4me1, which is correlated with a decrease in chromatin accessibility (53). BRG1 is also known to mediate inter-chromosomal looping interactions between specific loci such as the *MYC* enhancer and promoter, the alpha-globulin genes, the *IgH* locus, and the class II major histocompatibility complex gene locus (24, 61–64). On a global scale, BRG1 binding has been found at DNA-loop anchors (56) and topological associated domain (TAD) boundaries where it increases their stability (65). Together, this demonstrates an important role for BRG1 in maintaining chromatin architecture at both local and global levels for transcription regulation.

Here we dissected the molecular role of BRG1 on the transcriptome in prostate cancer. We confirmed that *SMARCA4* is over-expressed in prostate cancer irrespective of severity or cancer subtype and identified *SMARCA4* was also over expressed in a panel of prostate cancer cell lines. Depletion of BRG1 in LNCaP prostate cancer cells resulted in a modest effect on global gene transcription with most changes resulting in down-regulated gene expression. Within the cohort of down-regulated genes in BRG1 depleted cells we identified gene clusters defined by their co-occupancy or independence from transcription factors AR and FOXA1, both of which are known BRG1 co-activators (66–68). Our data revealed that BRG1, AR and FOXA1 co-regulate known prostate cancer genes *KLK2*, *PCAT1* and *VAV3*. Gene ontology analysis further revealed that genes regulated by BRG1 independent of AR and FOXA1 include factors regulating cell cycle and proliferation processes including DNA replication. In agreement, depleting BRG1 promoted G1 arrest resulting in reduced cell proliferation. Cumulatively the data indicate BRG1 promotes expression of cellular proliferation factors and cancer-associated genes in prostate cancer cells.

## RESULTS

### *SMARCA4* is over expressed in prostate cancer irrespective of tumour grade or subtype

We first examined the expression of *SMARCA4* in the TCGA (50) prostate normal and cancer cohort. The 486 tumour samples were subset into the seven TCGA categorised molecular subtypes of prostate cancer (50). These included those with fusion genes involving *ERG* (46%), *ETV1* (8%), *ETV4* (4%) and *FLI1* (1%), or those with mutations in *SPOP* (11%), *FOXA1* (3%) or *IDH1* (1%) (50). The remaining samples were grouped as ‘other’ (26%). Each subtype exhibited a statistically significant increase in *SMARCA4* expression (*p<0.05*) with the exception of the ‘FLI1’ subtype (*p=0.5899*) and ‘other’ (*p=0.1899*), which both demonstrated a non-significant increase in *SMARCA4* expression (Figure 1A). Previous work examining *SMARCA4* expression in the TCGA prostate cancer cohort demonstrated that it is also up-regulated irrespective of Gleason score (49). Therefore, we conclude that at the mRNA level, *SMARCA4* is universally over-expressed in prostate cancer, regardless of clinical grade or molecular subtype.

**Figure 1.**
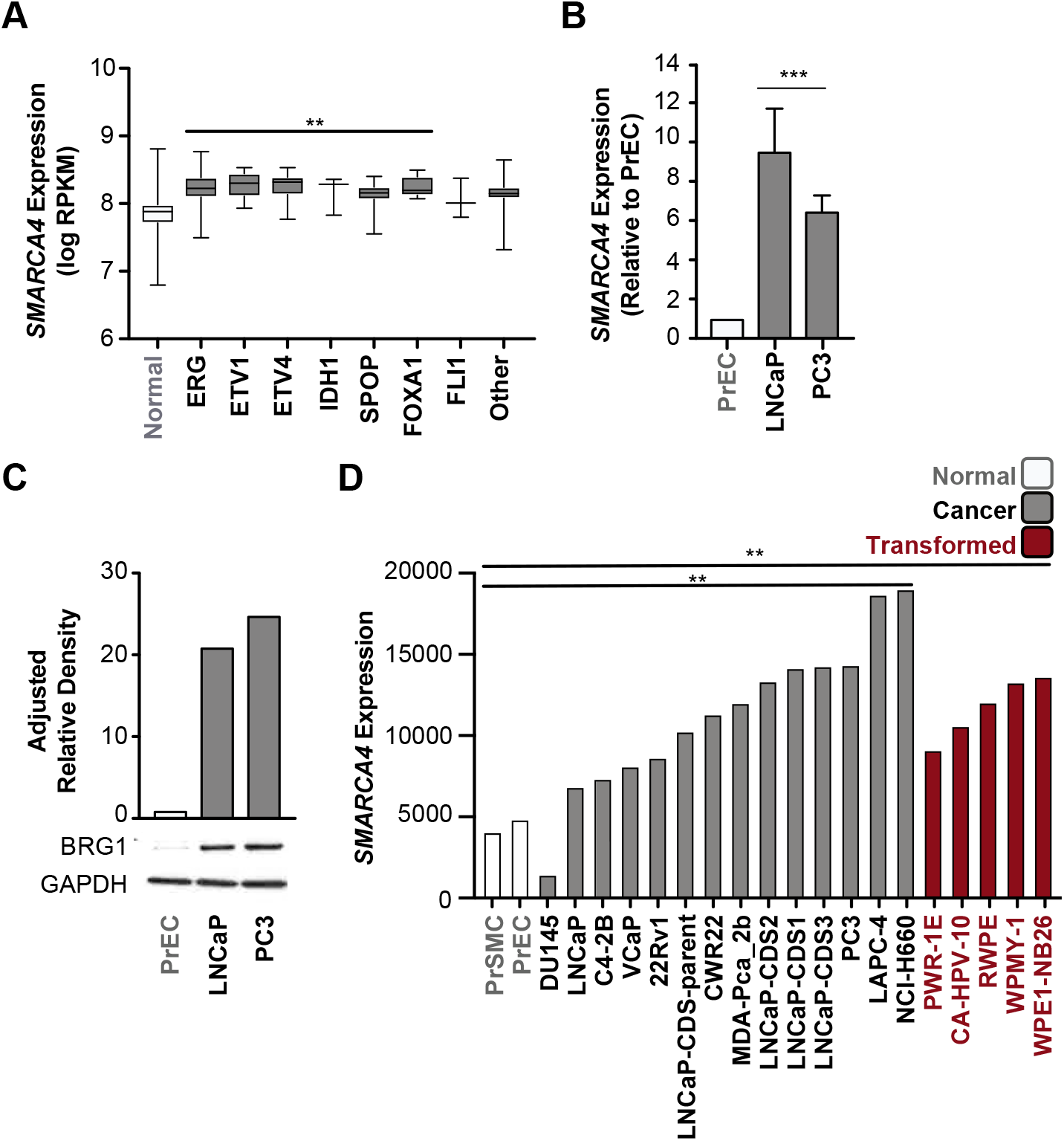
*SMARCA4* (BRG1) is over expressed in prostate cancer. A) *SMARCA4* gene expression (logPRKM) in TCGA data (tumours n= 486, normal = 52) with tumour samples separated by molecular subtype defined by the TCGA. *SMARCA4* expression is increased across all groups, with subtypes ERG, ETV1, ETV4, IDH1, SPOP, and FOXA1 all significantly up regulated, one-way ANOVA Dunnett’s multiple comparison correction \*\**p<0.05*. B) *SMARCA4* gene expression in prostate cell lines normalised to *18S* and relative to PrEC (n = 2). Significance determined by one-way ANOVA with Tukey’s multiple comparison correction \*\*\**p<0.001*. Bars denote mean, and error bars are SD. C) Representative Western blot of BRG1 protein level in prostate cell lines. Quantification above Western Blot by adjusted relative density normalized to GAPDH and relative to PrEC. D) Expression of *SMARCA4* from RNA-seq in prostate cell lines grouped as normal, cancer or transformed. The mean of each group was calculated, and a significance was tested by one-way ANOVA Dunnett’s multiple comparison correction, \*\**p<0.05*.

### *SMARCA4* is over expressed in prostate cancer and transformed prostate cell lines

We next examined both BRG1 protein and *SMARCA4* gene expression levels in normal prostate epithelial cells (PrEC) and compared to LNCaP (lymph node metastasis), an androgen-dependent prostate cancer cell line, as well as PC3 (bone metastasis), an androgen-independent prostate cancer cell line. We found that *SMARCA4* gene expression was increased ~9 fold in LNCaP cells and ~6 fold in PC3 compared to PrEC (*p<0.001*; Figure 1B). Further, the BRG1 protein level was increased ~20 and ~24 fold, respectively, in each of the prostate cancer cell lines compared to PrEC (Figure 1C). We compared this to published RNA-seq data of several normal, cancer and transformed prostate cell lines (69). The mean expression of *SMARCA4* was significantly increased in both the cancer cell lines and the transformed cell lines compared to the normal cells (*p=0.0148* and *p=0.0353* respectively; Figure 1D). The exception was DU145 cells that has a known frameshift mutation in *SMARCA4,* resulting in reduced expression (36). This data show that common prostate cancer cell lines reflect the same pattern of increased BRG1 protein that is observed in prostate tumours compared to normal prostate samples and therefore provides an appropriate model system to explore the functional consequences of BRG1 dysregulation on the transcriptome.

### BRG1 is required for the maintenance of active gene expression

Our previous work has shown that BRG1 occupancy is enriched at active promoter and enhancer gene regulatory elements in LNCaP cells (56). We therefore hypothesised that BRG1 would play an important role in maintaining the transcriptional profile of these cells. To assess this, we depleted the level of BRG1 protein using two independent siRNAs targeting *SMARCA4* (si-*SMARCA4*-1 and si-*SMARCA4*-2) and performed RNA-seq at 72 and 144 hours post transfection (Figure 2A). Initial assessment of our RNA-seq data confirmed successful depletion of the *SMARCA4* transcript (~80%) at both time points (Figure 2B). Additionally, we confirmed substantial depletion of BRG1 levels reduced to ~40% of the non-targeting control at 72 hours, and to ~20% of the non-targeting control at 144 hours post-transfection (Figure 2C). We note there were no significant changes detected in the gene expression of any other SWI/SNF subunit proteins (Supplementary Figure 1A). Further quality assessment of the RNA-seq data through a principal component analysis demonstrated that the samples separated by time-point on the first dimension, accounting for 43.39 % of the sample variance (Supplementary Figure 1B). We performed a differential gene expression analysis and identified 169 down-regulated genes and 24 up-regulated genes (logFC > 1.5, FDR < 0.05) at 72 hours post BRG1 depletion (Figure 2D). This increased to 800 down-regulated genes and 174 up-regulated genes by 144 hours post-transfection (Figure 2E). This suggests that the primary role of BRG1 in LNCaP cells is to maintain active gene expression of a subset of genes.

**Figure 2.**
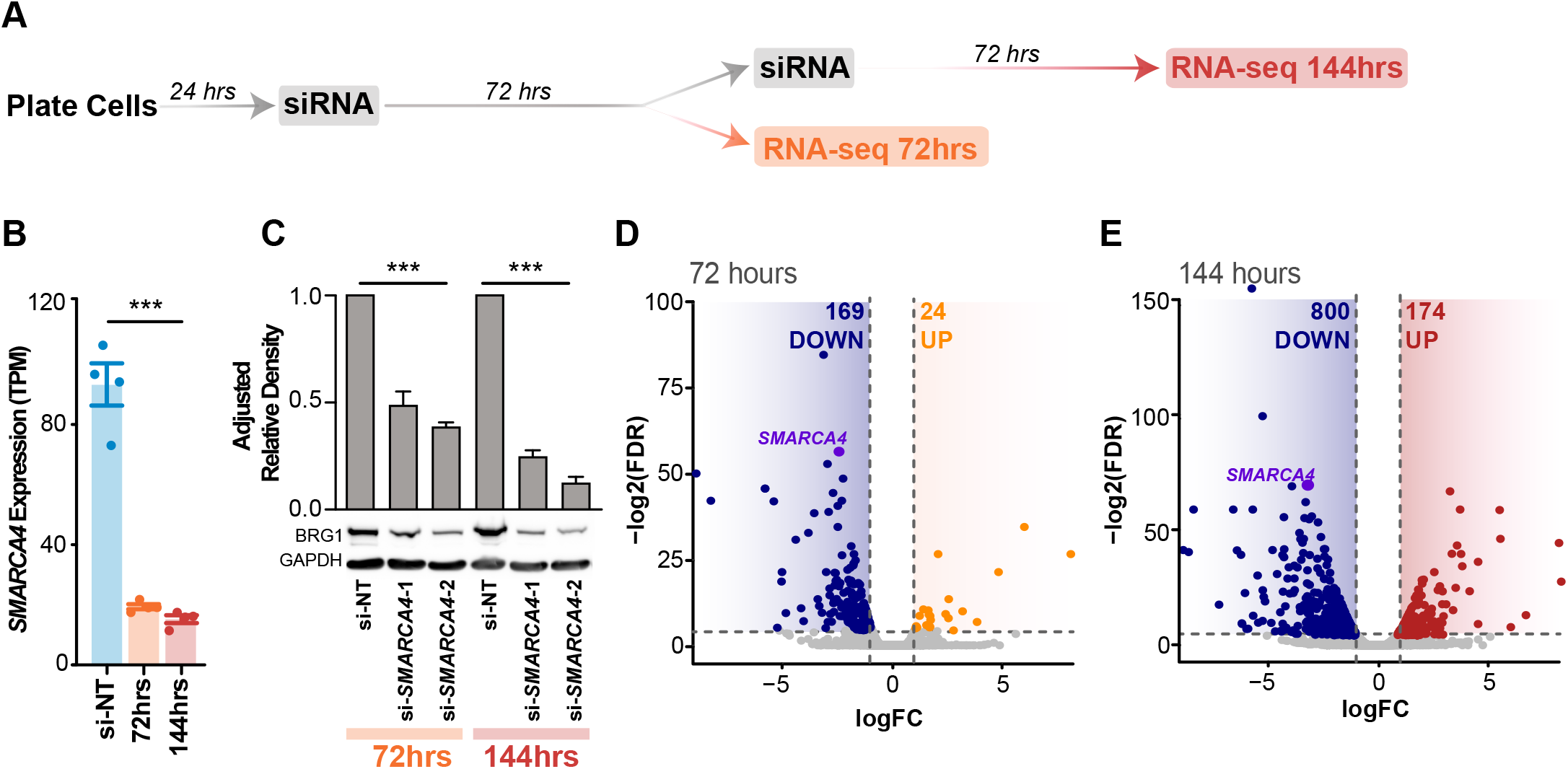
Loss of BRG1 results in a down regulation of gene expression. A) Schematic of temporal BRG1 knockdown model used for RNA-seq. Samples were collected at 72hrs (si-NT control, si-*SMARCA4*-1 and si-*SMARCA4*-2) and 144hrs (si-NT, si-*SMARCA4*-1 and si-*SMARCA4*-2) post siRNA transfection in duplicate for each condition at each time point (n=2). Cells were transfected with either control siRNA (si-NT) or *SMARCA4* siRNA. B) *SMARCA4* gene expression in control and post BRG1 depletion in the RNA-seq data, shown as transcripts per million reads (TPM). Control siRNA for 72 and 144 hours are shown collectively as si-NT. *SMARCA4* expression is significantly down regulated at both time points, *\*\*\*p<0.0001*. Bars denote mean, and error bars are SD. C) Representative Western blots of BRG1 and GAPDH protein levels at 72 and 144 hours post transfection. Adjusted relative density for BRG1 is calculated relative to GAPDH and normalized to the non-targeting control. Bars denote mean, and error bars are SD. D-E) Volcano plots of differentially expressed genes at 72 hours and 144 hours post knockdown. Significantly down regulated genes are blue and significantly up regulated genes for 72 and 144 hours post knockdown are shown in orange and red respectively. *SMARCA4* differential expression is highlighted in purple. Expression is shown as normalised log2 counts per million reads.

### BRG1 does not function in the regulation of alternative splicing

The nucleosome barrier within genes is reported to contribute to alternative splicing, where there is a higher conservation of nucleosomes at the splice sites of constitutive exons compared to skipped exons (70–72). Since the contribution of BRG1 to alternative splicing regulation is unknown, we investigated if alterations in alternative splicing may explain down regulation of gene expression after BRG1 depletion in LNCaP cells. To do this we performed a multivariate analysis of transcript splicing (MATS; (73–75)) of our RNA-seq datasets. After 72 hours of BRG1 depletion, MATS pairwise comparison detected a genome wide total of 13 and 11 skipped exons, and 14 and 9 retained introns with si-*SMARCA4*-1 and si-*SMARCA4*-2 respectively (Supplementary Figure 1C). At 144 hours post BRG1 knockdown this increased to 240 and 260 skipped exons, and 27 and 26 retained introns with si-*SMARCA4*-1 and si-*SMARCA4*-2, respectively (Supplementary Figure 1D). Given the relatively large number of intron-exon junctions within the total LNCaP transcriptome, we conclude BRG1 does not extensively contribute to alternative splicing as the mechanism for predominant gene down-regulation. However, we do note that at 144 hours post-knockdown the MATS analysis identified retention of the first intron from the Kallikrein 3 gene, which encodes prostate specific antigen (PSA) (Supplementary Figure 1E). This splice variant has previously been reported in LNCaP cells and generates a unique protein from canonical PSA (76). While PSA has a well-known link to prostate cancer, the function of its alternative splice variant remains unknown.

### BRG1 binding is associated with expression of prostate cancer associated genes

We further examined our RNA-seq datasets to determine which genes showed a significant change in expression at 72 hours that was maintained at 144 hours. Of the genes that were down-regulated at the 72 hour time point, 126 genes (75 %) remained down-regulated at 144 hours. Similarly, of the up-regulated genes, 16 (67 %) remained up-regulated at the extended time point (Figure 3A). Within the down-regulated gene set we note a number of genes that have previously been associated with increased proliferation in prostate cancer; these include kallikrein 2 (*KLK2*), long non-coding RNA prostate cancer associated transcript 1 (*PCAT1*), Vav guanine nucleotide exchange factor 3 (*VAV3*) (69, 77–84) (Figure 3B-D). We also examined the panel of prostate cell lines (69) and confirmed that, on average there is elevated expression of these genes in both prostate cancer cells and transformed prostate cell lines compared to normal prostate cells (Supplementary Figure 2A). This suggests a role for BRG1 in maintaining the expression of genes associated with prostate cancer proliferation.

**Figure 3.**
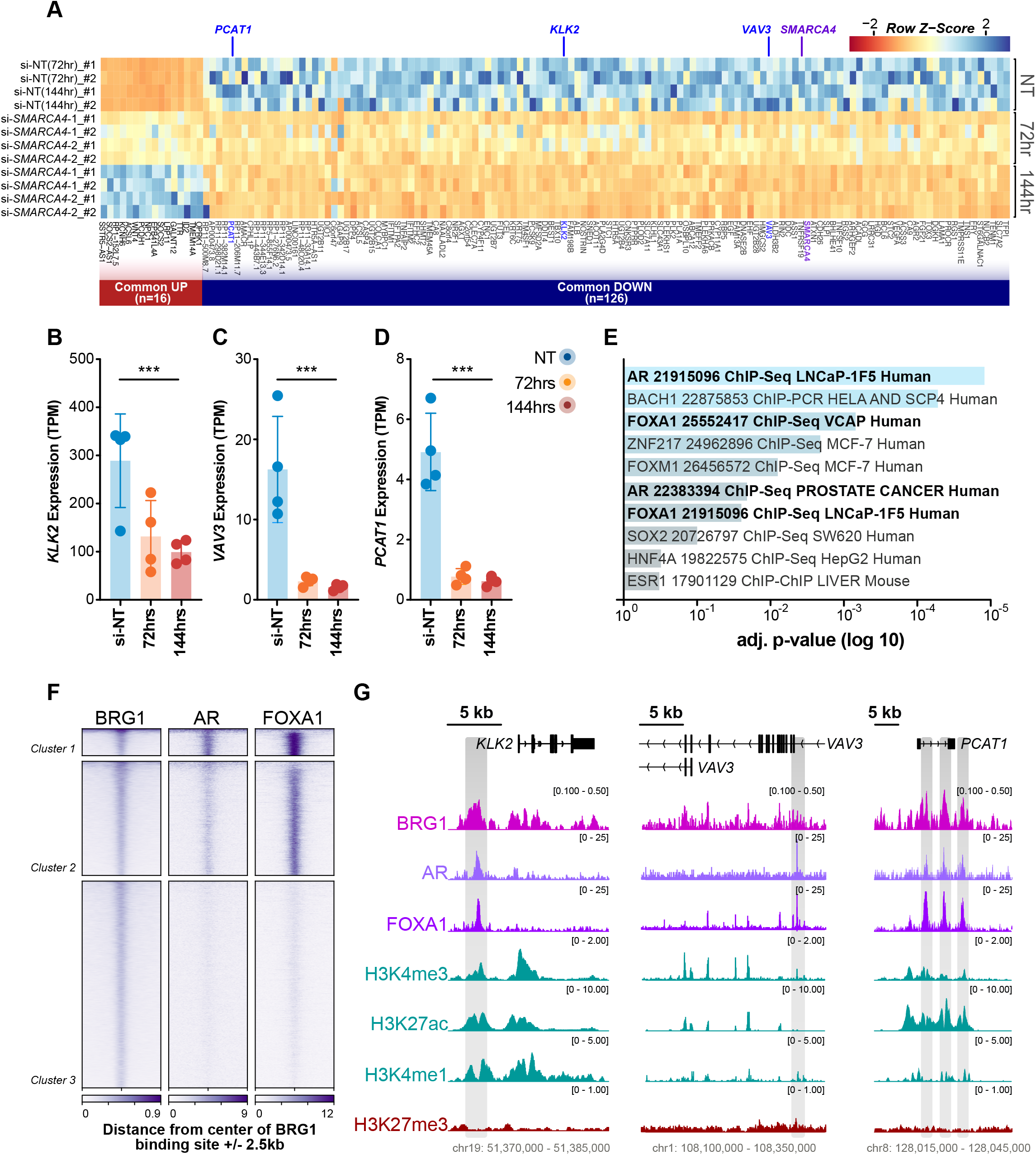
BRG1 regulates genes associated with prostate cancer. A) Heatmap illustrating RNA-seq differential gene expression data for up (n = 16) and down (n = 126) regulated genes common to both time points after BRG1 depletion. Expression is represented as the normalised row Z-score of TPM. B-D) *KLK2, VAV3 and PCAT-1* gene expression from the RNA-seq datasets shown as TPM. Bars denote mean, and error bars are SD. E) Gene set enrichment analysis using ‘Enrichr’ of differentially expressed genes that are common to both time points, showing the adjusted p-value (log 10, reversed x-axis) of significantly enriched transcription factor ChIP-seq from ChEA curated data *(p<0.05)*. F) Heatmap of BRG1, AR and FOXA1 ChIP-seq signal at BRG1 binding sites in LNCaP cells, +/− 2.5 kb from the centre of the binding site. Data is clustered into three groups by k-means. G) IGV images of the genes *KLK2*, *PCAT-1* and *VAV3*. Grey shaded regions contain ChIP-seq signal peaks for BRG1, AR and FOXA1.

We next sought to further explore commonalities in the genes with a significant change in expression at both time points. We used ‘Enrichr’ (85, 86) to determine which existing ChIP-seq datasets of transcription factors had enriched binding at the promoters of these genes. We discovered that the most significantly enriched datasets were for the androgen receptor (AR) and Forkhead box A1 (FOXA1) (Figure 3E), both of which are important for prostate cancer growth (66, 67, 87–91). To investigate the potential coordinated function of these transcription factors with BRG1, we compare the ChIP-seq signal of BRG1 (91), AR (87) and FOXA1 (87) at BRG1 genome-wide binding sites in LNCaP cells. We found the profiles separated into three clusters. Cluster 1 sites displayed strong AR and FOXA1 binding, cluster 2 had moderate AR and strong FOXA1, and cluster 3 had minimal to no signal for AR or FOXA1 (Figure 3F). We next examined the key BRG1 regulated genes *KLK2*, *PCAT1* and *VAV3*, and found coordinated binding of all three factors at the promoters of *KLK2* and *PCAT1*, and binding of BRG1 and FOXA1 upstream of the internal 3-prime promoter of *VAV3* (Figure 3G). Furthermore, we showed that the expression of AR or FOXA1 themselves was not regulated by BRG1 (Supplementary Figure 2B-C), suggesting that the loss of BRG1 is enough to disrupt expression and regulation of *KLK2*, *PCAT1* and *VAV3*.

### BRG1 binding is associated with the expression of DNA replication genes

As the majority of significant gene changes occurred at 144 hours post-knockdown, we next investigated potential gene regulatory networks. Gene ontology analysis with Enrichr (85, 86) identified several significant (FDR < 0.05) GO terms pertaining to biological processes, cellular component and molecular function that were all broadly related to the cell cycle (Figure 4A). As BRG1 has previously been shown to interact with cell cycle master regulators, such as Rb and p53 (19, 92–94), we explored the relationship between the cell cycle and BRG1 further in our datasets. We compiled a list of 250 genes related to cell cycle processes, curated from the cell cycle GO terms, and of these examined the top 40 most significantly down-regulated genes in our dataset. Of note among the list were several key genes involved in DNA replication initiation such as *CDC6*, *CDT1* and *CDC45*, as well as the Minichromosome Maintenance (MCM) replicative helicase components *MCM2* and *MCM5* (Figure 4B). To investigate if the effect on replication initiation gene expression was more widespread, we reviewed the gene expression of the other components in the MCM2-7 replicative helicase and the Origin Recognition Complex (ORC) and found that several of these genes were also down-regulated (Figure 4C-D). We confirmed the down regulation of MCM5, CDC6 and ORC6 via Western blot, along with cell cycle regulator CHK1, which revealed almost undetectable expression by 144 hours post BRG1 knockdown (Figure 4E-F).

**Figure 4.**
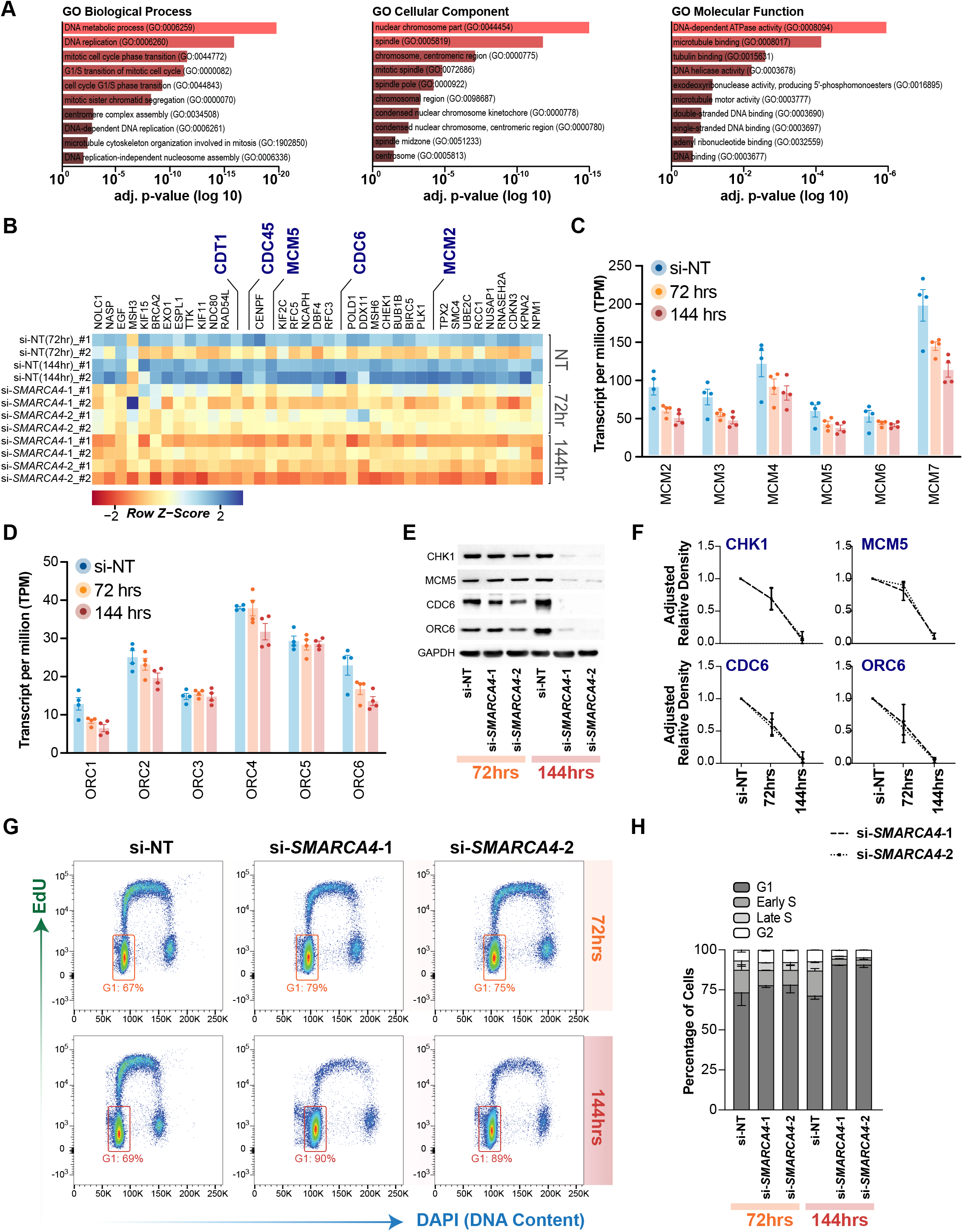
BRG1 regulates genes involved the cell cycle. A) Gene set enrichment analysis using ‘Enrichr’ of down regulated genes at 144 hours post BRG1 knockdown. Enriched GO terms are classified as biological processes, cellular component or molecular function. Adjusted p-value (log 10, reversed x-axis) of the 10 most significant GO terms are shown. B) Heatmap of gene expression profiles from the top 40 differential cell cycle genes after BRG1 depletion. Expression is shown as the normalised row Z-score of transcripts per million reads (TPM), with blue indicating higher expression and red indicating lower expression. Genes involved in DNA replication initiation are indicated in blue. C-D) Gene expression from RNA-seq (TPM), of the MCM2-7 helicase components (top) and the Origin of Replication complex (ORC) subunits (bottom). Error bars denote mean and standard deviation. Bars denote mean, and error bars are SD. E) Representative Western blot showing protein levels of replication initiation genes MCM5, CDC6 and ORC6, along with CHK1, after 72 and 144 hours post BRG1 depletion. Error bars demonstrate SD. F) Quantification of Western blots demonstrating adjusted relative density to GAPDH (n = 2). G) Representative flow cytometry scatter of DAPI (x-axis) and EdU (y-axis) fluorescence intensity at 72 and 144 hours post BRG1 knockdown. G1 cells are shown by boxed gate. H) Percentage of cells in each phase of the cell cycle from flow cytometry data, error bars show standard deviation (n = 2). Error bars show SD.

We investigated whether AR and FOXA1 were also colocalised with BRG1 at DNA replication genes. We examined the ChIP-seq binding profiles of AR, FOXA1 and BRG1 at the promoters of 91 DNA replication genes (determined from the DNA replication GO terms) that were expressed in LNCaP cells. We found at promoters of these genes containing the active histone marks H3K4me3 and H3K27ac, also displayed a weak BRG1 ChIP-seq signal, but were completely absent of AR and FOXA1 ChIP-seq peaks (Supplementary Figure 3A), for example at the promoters of *CDC45*, *ORC6* (Supplementary Figure 3B). Additionally, we also note this pattern at a putative enhancer region within the *MCM2* gene (Supplementary Figure 3B). Our data suggests that BRG1 binding is associated with the expression of DNA replication genes in prostate cancer cells that is independent of AR and FOXA1.

### BRG1 depletion arrests cells in G1

Given BRG1 regulates several genes involved in proliferation and replication, we next asked if BRG1 depletion would alter cell cycle progression in LNCaP cells. We investigated this utilising the same siRNA-mediated approach to target BRG1 by depleting *SMARCA4* and conducted flow cytometry cell cycle analysis at 72 and 144 hours post knockdown. We detected an increase of cells in G1 at 72 hours, which was enhanced by 144 hours. Specifically, at 144 hours post BRG1 depletion there was ~20% increase of cells in G1 and equivalent loss of cells in S phase (Figure 4G-H). These data suggest that a loss of BRG1 reduces proliferation through mediating a G1 arrest.

## DISCUSSION

Here we examined the involvement of the SWI/SNF chromatin remodeller BRG1 and its associated encoding gene *SMARCA4* in prostate cancer transcriptional deregulation. We found that over expression of *SMARCA4* commonly occurs in both the TCGA prostate cancer cohort, irrespective of tumour subtype, and in a panel of prostate cancer cell lines. We also found that knockdown of the *SMARCA4* gene, and consequently the BRG1 protein, results in down-regulation of pro-proliferative transcriptional pathways. These included genes already known to promote prostate cancer proliferation, as well as cell cycle and DNA replication genes. Reduction of gene expression in these pathways was concomitant with G1 arrest. Taken together, our results provide new insights into BRG1’s contribution to transcriptional patterns relating to proliferation in prostate cancer.

We have demonstrated that *SMARCA4* mRNA over expression is a universal feature of prostate cancer. Clinical datasets have shown BRG1 protein levels are over-expressed in prostate cancer, in the absence of consistent significant deleterious genetic mutations evident in *SMARCA4* (42, 44–46). Using the large prostate cancer cohort from TCGA (50) we found that *SMARCA4* was significantly over-expressed. Consistent with this, *SMARCA4* expression was increased in a panel of both prostate cancer and transformed cell lines. These data emphasise that the overall increased expression of *SMARCA4* is a characteristic of prostate cancer, irrespective of subtype.

BRG1 depletion followed by RNA-seq revealed multiple transcriptomic alterations that were regulated by BRG1 and related to proliferation. BRG1 depletion primarily resulted in the down-regulation of BRG1’s target genes, indicating the main role of BRG1 is to promote active gene expression. Within the down-regulated genes were genes associated with increased proliferation in prostate cancer including *KLK2*, *PCAT-1* and *VAV3*. KLK2 is a known activator of PSA, which is an important biomarker of prostate cancer, and associated with decreased apoptosis (77, 84). *PCAT-1* promotes proliferation through the oncoprotein Myc (69, 81), while VAV3 regulates AR activity to stimulate growth in prostate cancer (78–80, 82). Both *PCAT-1* and VAV3 are correlated with disease progression. Through an analysis of gene ontologies, we also found several cell cycle gene pathways were downregulated with BRG1 depletion. This included numerous genes involved in DNA replication, which were among the most significantly down regulated genes following BRG1 depletion. BRG1 is known to have a role in driving self-renewal and malignancy in B-cell acute lymphoblastic and acute myeloid leukaemias, cancers which also have over expressed BRG1 (22, 24). Specifically, these leukaemias require high levels of BRG1 for de-condensation of the cell specific *MYC* enhancer. In these cancers, a loss of BRG1 causes a reduction of enhancer-promoter interactions, reduced transcription factor occupancy and DNA looping which in turn reduces *MYC* expression (24). This implies that the overexpression of BRG1 contributes to driving oncogenic transcriptional programs which influence the proliferation capacity of cancer cells.

Our data revealed that BRG1 co-occupied the promotors of proliferation associated genes (*KLK2*, *PCAT-1* and *VAV3)* along with AR and FOXA1, and that these genes were down-regulated across our experimental time course. Co-regulation of transcription by AR and FOXA1 in prostate cancer is associated with reprogrammed binding of AR and oncogenic patterns of gene expression that are essential for AR-driven proliferation (95, 96). Additionally, there is a high overlap of these reprogrammed AR binding sites between LNCaP cells and primary prostate tumour tissue (96). Here we have shown BRG1 gene regulation overlaps with these transcription factors at gene promoters, which is concomitant with expression of prostate cancer associated genes. However, it is noteworthy that BRG1 depletion also altered the expression of DNA replication genes through a mechanism that appears independent of AR and FOXA1. This data suggests that BRG1 has additional roles in other gene regulatory networks, which may indirectly influence cell proliferation. As BRG1 is known to interact with cell cycle regulators in other cancers, it is possible that genes co-regulated by BRG1, AR and FOXA1 are important in a prostate cancer context, while regulation of cell cycle and DNA replication genes may be a general feature of BRG1 over expression in cancer.

## CONCLUSIONS

In summary, our data identifies fundamental role for BRG1 in maintaining active transcription for proliferation of prostate cancer cells. We find that BRG1 promotes gene expression in prostate cancer models with varying degrees of dependence on AR and FOXA1. BRG1 is required to drive the expression of numerous prostate cancer specific genes in an AR/FOXA1 dependant manner, but also works independently to drive the expression of pro-proliferative and DNA replication genes. These results provide important functional information regarding the role of BRG1 controlling proliferation in prostate cancer cells.

## METHODS

### Cell Culture and siRNA Transfection

Normal Prostate Epithelial Cells (PrEC) cells (Cambrex Bio Science, #CC-2555) were cultured in PrEBM (Clonetics, #CC-3165) according to the manufacturer’s protocol. Briefly, PrEC cells were seeded at 2,500 cells per cm^2^ and medium was replaced every two days. Cells were passaged at approximately 80 % confluence. To passage a T75 flask, PrEC cells were rinsed in 6 mls Hanks Balanced Salt Solution (Thermo Fisher Scientific, #14025076) then detached with 2 ml pre-warmed 0.025 % Trypsin-EDTA and incubated at room temperature for 5 minutes. Trypsin was inactivated with 12 mls of Trypsin-Neutralizing Solution (Clonetics, #CC-5002) and cells were centrifuged at 300 x g for 5 minutes. The supernatant was aspirated, and the cell pellet was re-suspended in PrEBM. The number of cells was determined on the Countess automated counter and were re-seeded at the appropriate density based on experimental needs. Cells were discarded after ~16 population doublings.

PC3 cells (ATCC, #CRL-1435) were maintained in RPMI medium (Gibco, #11875-093) with 10 % FBS, 11 mls of 1 M HEPES (Gibco, #15630080) and Pen/Strep. LNCaP cells (ATCC, #CRL-1740) were cultured using custom T-Medium from Gibco (DMEM low glucose (GIBCO cat# 31600-034), Kaighn’s modified F-12 medium (F-12K; cat# 211227-014), insulin 500x bovine pancreas (Sigma cat# I1882.10MG), T3 6.825 ng/ml Tri-iodothyronine (Sigma cat# T5516), Transferrin 500x (Sigma cat# T5391), Biotin 500x (Sigma cat# B4639), Adenine 500x (Sigma cat# A3259)). Both prostate cell lines were cultured under recommend conditions; 37°C with 5 % CO_2_. When the cells reached ~80 % confluence they were passaged or seeded as per experimental requirements. For siRNA transfection LNCaP cells were seeded into 6-well plates at a density of 2.5 × 10^5^ cells per well or 10cm dishes at 1.5 × 10^6^ cells per dish. The cells were transfected with either on target *SMARCA4* siRNA (Horizon, #J-010431-06-0005 or #J-010431-07-0005) or the non-targeting control siRNA pool (Horizon, #D-001810-10-05) 24 hours after seeding the cells using DharmaFECT 2 (Thermo Scientific, #T-2002-03) as per the manufacturer’s instructions. To maintain the knockdown over a 6-day period, at 72 hours post transfection the cells were harvested, split at a ratio of 1:2 into two new wells, and reverse-transfected with siRNA. The cells were then incubated for a further 72 hours before collection.

### Quantitative Real-Time PCR (qRT-PCR)

RNA was extracted with TRIzol reagent (Thermo Scientific, #15596026), according to the manufacturer’s protocol. Extracted RNA was re-suspended in 30 ul of nuclease-free water and quantified on the NanoDrop spectrophotometer (Thermo Scientific). cDNA synthesis was carried out with 500 ng of RNA using the SensiFAST cDNA Synthesis Kit (Bioline, #BIO-65054) according to the manufacturer’s instructions.

qRT-PCR was carried out on the CFX384 Touch Real-Time PCR Detection System (Bio-Rad). A master mix was made for each qRT-PCR target containing 5 ul of KAPA Universal SYBR Fast PCR mix (KAPA Biosystems, #KK4602), 0.6ul of 5 uM forward primer, 0.6 ul of 5 uM reverse primer and 1.8 ul of nuclease free water per reaction. Reactions conditions were 95°C for 3 minutes, followed by 45x cycles of 95 °C for 3 seconds and 60 °C for 30 seconds, then a melt curve analysis (65 °C to 95 °C, increasing at a rate of 0.5 °C every 5secs). Primers to detect *SMARCA4* were CAGAACGCACAGACCTTCAA (forward) and TCACTCTCCTCGCCTTCACT (reverse) and for detection of *18S* GGGACTTAATCAACGCAAGC (forward) and GCAATTATTCCCCATGAACG (reverse). Relative gene expression was calculated using ddCt and normalised to *18S*. A significant change in gene expression of *SMARCA4* between PrEC, LNCaP and PC3 cells was determined by one-way ANOVA and corrected with Tukey’s test for multiple comparisons.

### Western Blot

Whole cell lysates were collected with lysis buffer (50 mM HEPES, 150 mM NaCl, 10% Glycerol, 1 % Triton-X-100, 1.5 mM MgCl_2_, 1 mM EGTA, 10 mM Pyrophosphate, 100 mM NaF, Roche protease inhibitor cocktail 1x), and protein level quantified using the Pierce BCA Assay Kit (Thermo Scientific, #23227) according to the manufacturer’s instructions. Sample reducing agent (Thermo Scientific, NP0004), loading buffer (Thermo Scientific, NP0007) and 10 ug protein were combined with water to a final volume of 25 ul. Protein samples were heated at 90 °C for 5 minutes then allowed to cool to room temperature. Protein samples were loaded on a NuPage Novex Bis-Tris 4-12 % gel (Thermo Scientific, NP0321BOX) and electrophoresed at 100V for 1.5 hours in a 1x MOPS buffer (50 mM MOPS (Biochemicals Astral Scientific, #BIOMB03600, 50 mM Tris base, 0.1% SDS, 1 mM EDTA [pH 7.7]). Proteins were transferred to a polyvinylidene fluoride membrane (Bio-Rad, #1620177) at 30 volts for 1 hour using 1x transfer buffer (25 mM Tris base, 192 mM Glycine [pH 8.3]) with 10 % methanol (Sigma-Aldrich, #322415). Membranes were blocked for 1 hour with 5 % skim milk in TBS-T (20 mM Tris, 150 mM NaCl, 0.1% Tween 20 [pH 7.6]) at 4°C. Primary antibodies used were BRG1 (Santa Cruz, sc-10768X), GAPDH (Ambion, AM4300), CHK1 (CST, 2360S), ORC6 (CST, 4737S), CDC6 (CST, 3387S) and MCM5 (abcam, ab17967). Primary antibodies were incubated on samples overnight at 4 °C with rotation. The membrane was then washed three times for 10mins each in TBS-T with rotation. Secondary antibodies goat anti-mouse (Santa Cruz, sc-2005) and goat anti-rabbit (Santa Cruz, sc-2004) were diluted in TBS-T containing 5 % skim milk and incubated at 4 °C with rotation for 1 hour. The membrane was washed three times for 10 minutes in TBS-T. The membrane was then covered with ECL solution (Perkin Elmer, #NEL104001EA), incubated for 1min at room temperature, and visualized by X-ray film. Adjusted relative density calculations were processed through ImageJ (97, 98).

### Flow cytometric cell cycle analysis

LNCaP cells were seeded at 1.5 × 10^6^ cells per 10 cm dish and transfected with siRNA as described. At 72 and 144 hours post transfection the cells were treated with 10 uM EdU for 30 minutes. Remaining EdU was washed off the cells with PBS before harvesting cells, then 1 × 10^6^ cells were fixed in 70 % ethanol and frozen at −20 °C. Cells were then diluted 1 in 4 with PBS then pelleted and re-suspended in 1 ml of PBS containing 1 % BSA (Sigma-Aldrich, #A2058). Cells were again pelleted, re-suspended in 500 ul of click reaction mix (10uM carbocyfluorescine TEG-azide, 10 mM Sodium L-ascorbate, and 2 mM Copper-II-sulphate diluted in PBS), and incubated in the dark at room temperature for 30 minutes. Samples were then diluted with 5 mls of PBS containing 1 % BSA and 0.1 % Tween-20. Cells were again pelleted, washed with PBS and then resuspended in 500 ul of PBS containing 1% BSA, 0.1 mg/ml of RNase and 1 ug/ml of DAPI. Samples were analyised on the Canto II (BD Biosciences). Forward and side scatter were used to select a population of cells free of cell debris and doublets. Cells were analysed using B450 (FTIC – EdU positive) and B510 (DAPI) lasers. 50,000 single cell events were recorded for each ample. FlowJo software v10.5, was used to analyse the data. Data was collected in biological duplicate.

### RNA-seq Experiments

Total RNA was extracted with TRIzol reagent, quantified on the Qubit and quality assessed with the Bioanalyzer. An aliquot of 500 ng of total RNA was spiked with external controls ERCC RNA spike-in Mix (Thermo Scientific, 4456740) and libraries constructed with the TruSeq Stranded mRNA sample preparation kit (Illumina, 20020594) according to the manufacturer’s protocol. mRNA Libraries were quantified on Qubit and then stored at −20 °C. Library quality and fragment size of RNA-seq libraries was assessed on the Bioanalyzer, then KAPA Library Quantification (KAPA Biosystems, #KK4824) was performed according to the manufacturer’s protocol. The KAPA quantification results were used to dilute the libraries to 2 nM for sequencing. RNA-seq samples were sequenced for 100 cycles of paired-end reads on the Illumina HiSeq 2500 platform, with four samples multiplexed per lane of the high output run.

### RNA-seq Data Analysis

RNA-seq data was processed as described in Taberlay & Achinger-Kawecka *et al.* (9). Briefly, read counts were normalized with ERCC spike in controls, mapped to hg19/GRCh37 using STAR and counted into genes using the featureCounts (99) program. GENCODE v19 was used as a reference transcriptome to determine the transcript per million read (TPM) value. Fold change was calculated within each time point as the log2 ratio of normalized reads per gene using the *edgeR* package in R. Genes with a fold change of ± 1.5 and FDR < 0.01 were considered significantly different. Volcano plots of differential expression were created in R with *ggplots2* and heatmaps with the *heatmap2* package with normalised row Z-score. PCA was performed in R using the *edgeR* package with log counts per million (logCPMS) over GENCODE v19 annotated gene coordinates and normalizing the read counts to library size. RNA-seq multivariate analysis of transcript splicing (MATS) to calculate exon skipping and intron retention was performed with the MATS python package v4.0.2 (73–75). Transcription factor and GO term enrichment was obtained from Enrichr (http://amp.pharm.mssm.edu/Enrichr/) online gene list analysis tool (85, 86).

### TCGA and prostate cell line expression analysis

Pre-processed RNA-seq data from the TCGA prostate adenocarcinoma cohort was downloaded (cancergenome.nih.gov) for both normal and tumour samples. The average of tumour (n = 486) and normal (n = 52) samples was calculated to determine mean expression. Separation of tumours by Gleason score and molecular subtype was performed in R using the associated clinical data to subset the appropriate groups. Significance was calculated for tumour versus normal using an unpaired T-test. For comparison between Gleason score or molecular subtype, significance was calculated using one-way ANOVA with Dunnett’s multiple comparison correction.

Expression data for prostate cell lines from Presner et al. (69) was downloaded from http://www.betastasis.com/prostate_cancer/. Significance between normal, cancer and transformed cell lines was calculated using one-way ANOVA with Dunnett’s multiple comparison correction.

### ChIP-seq data

The following LNCaP ChIP-seq data was obtained from GEO (ncbi.nlm.nih.gov/geo/); BRG1 accession GSE72690 (91), H3K4me3 and H3K27me3 accession GSE38685 (100), H3K27ac and H3K4me1 accession GSE73785 (9). These data were processed through NGSane pipeline as previously described (9, 100). Pre-processed bigwig files for FOXA1 and AR were obtained from GEO accession GSE114274 (87). Genome browser images of ChIP-seq data were taken from IGV. Heatmaps of ChIP-seq signal were created with *deeptools* (101).

## DECLARATIONS

### Ethics approval and consent to participate

Not Applicable

### Consent for publication

Not Applicable

### Availability of data and materials

The BRG1 knockdown RNA-seq data generated for this study has been submitted to GEO, accession number GSE150252. Reviewer access for the submitted data is available from; https://www.ncbi.nlm.nih.gov/geo/query/acc.cgi?acc=GSE150252

### Authors’ contributions

This study was initiated and deisgned by KAG, SJC and PCT. Experiments were performed by KAG, SGP and GK. Analysis and interpretation of next-generation data was performed by KAG, CMG, JAK and PL. Initial manuscript draft was written by KAG. Manuscript editing and reviewing was conducted by KAG, JAK, SGP, GK, AJC, SJC and PCT. All authors have read and approval the final version of this manuscript. Funding for this work was provided by AJC, SJC and PCT.

## Acknowledgements

We thank Suat Dervish and the Westmead Flow Cytometry facility for the FACS analysis infrastructure.

---

**Supplementary Figure 1.** A) SWI/SNF subunit gene expression (TPM) from RNA-seq data. All subunits, except *SMARCA4* (shown in Figure 2A), are not significantly altered. Bars denote mean, and error bars are SD. B) PCA plot characterising the trend in expression profiles between the non-targeting control and after BRG1 knockdown. Each point on the plot represents an RNA-seq sample. Samples are separated by principal components 1 and 2, which together explain 58.37 % of the variance between the samples. C) Number of skipped exons at 72 hours and 144 hours after BRG1 knockdown with si-*SMARCA4*-1 (black) and si-*SMARCA4*-2 (grey). D) Number of retained introns at 72 hours and 144 hours post BRG1 depletion with si-*SMARCA4*-1 (black) and si-*SMARCA4*-2 (grey). E) Sashimi plot of exons one and two of the *KLK3* gene in the non-targeting and 144 hour knockdown RNA-seq data. Arcs represent the number of split reads across the exons. Lower numbers represent increased retention of the first intron after BRG1 knockdown.

**Supplementary Figure 2.** A) Expression of *KLK2, PCAT-1 and VAV3* in prostate cell lines grouped as normal, cancer or transformed. B) *AR* and *FOXA1* gene expression from the RNA-seq datasets shown as TPM. Bars denote mean, and error bars are SD.

**Supplementary Figure 3.** A) Heatmap of replication gene promoters, +/− 5kb from the transcription start site. B) IGV images of the genes *CDC45*, *ORC6* and *MCM2*. Grey shaded regions contain ChIP-seq signal peaks for BRG1 and active histone modifications.

