## Supplementary figures and images for "BRG1 promotes transcriptional patterns that are permissive to proliferation in cancer cells"

### Supplementary Figures 1-3

#### SUPPLEMENTARY FIGURE 1

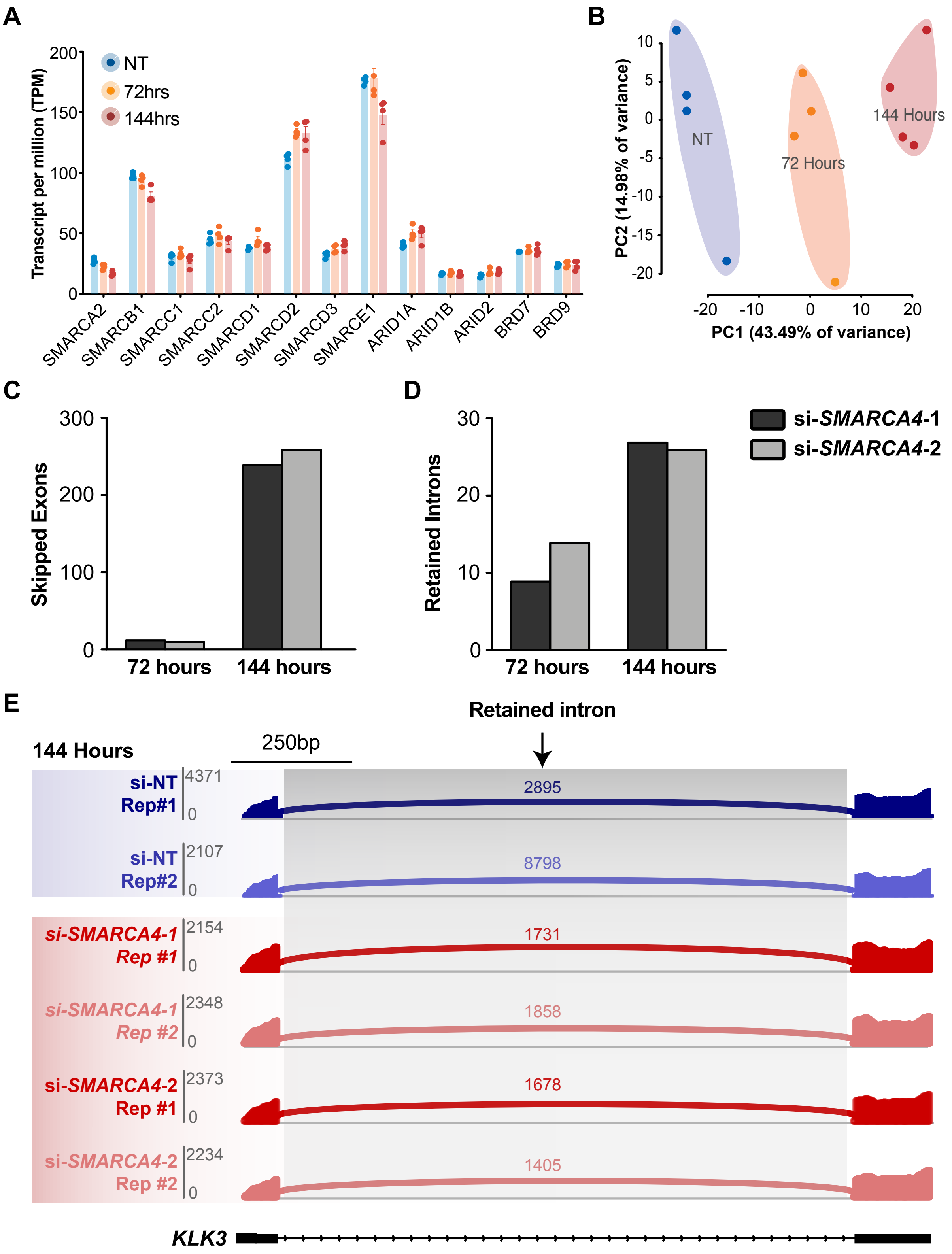

#### SUPPLEMENTARY FIGURE 2

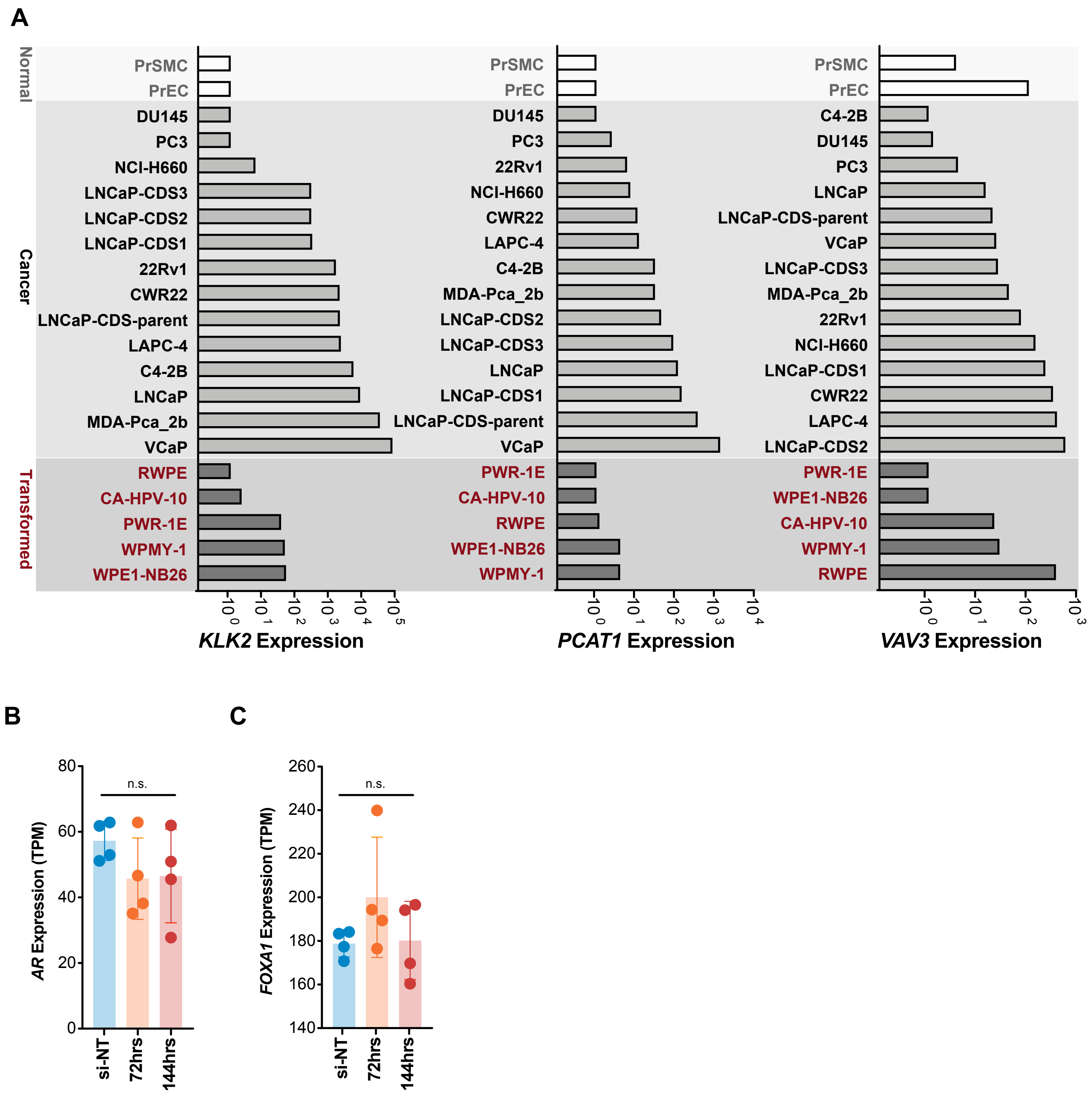

#### SUPPLEMENTARY FIGURE 3

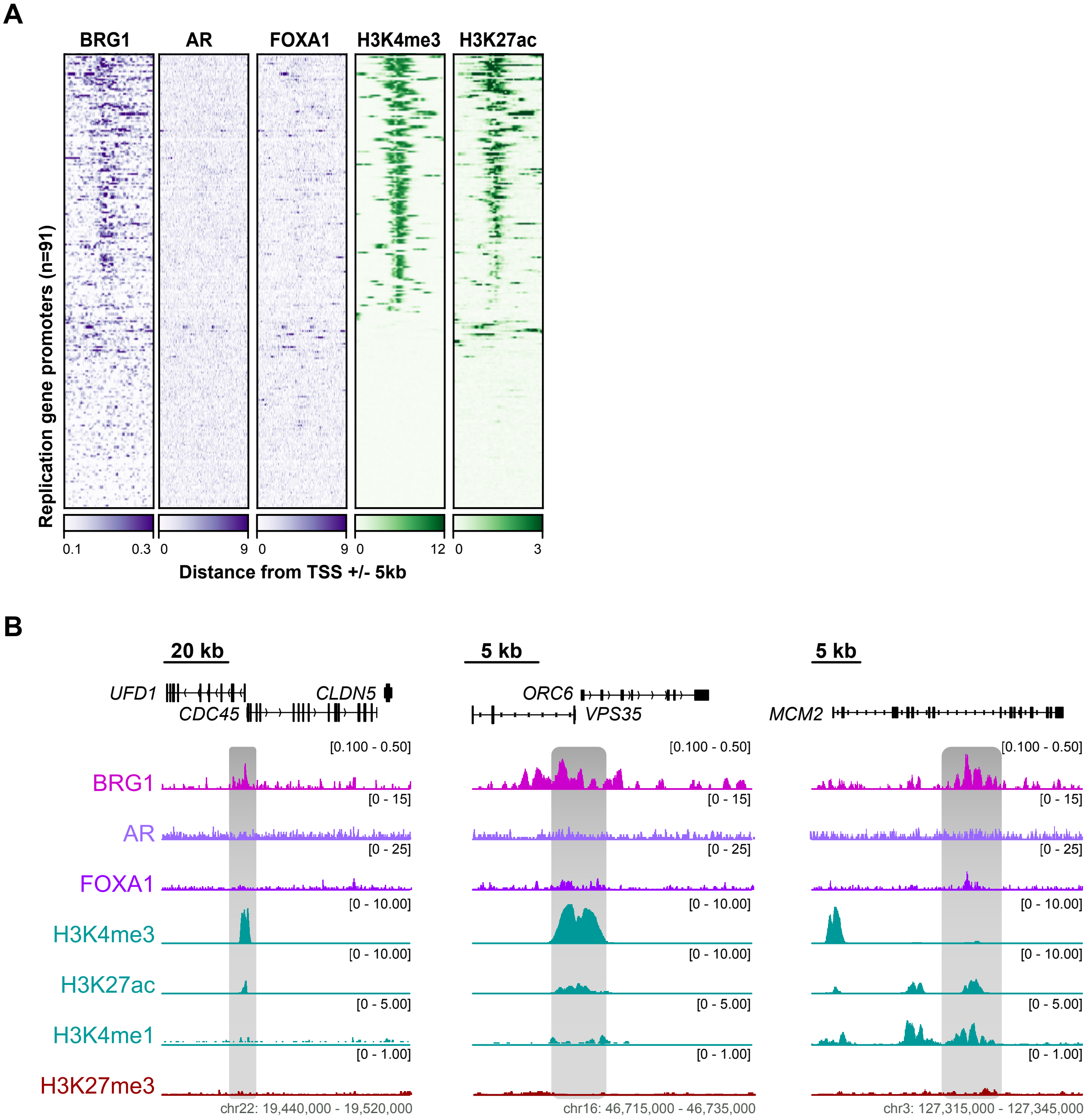
